# CD19-CAR T-cell therapy reshapes the host–microbiota immune axis in autoimmune diseases

**DOI:** 10.64898/2026.11.11.765834

**Authors:** Yuichi Maeda, Yi-Yuan Lee, Sylvio Redanz, Viacheslav Petrov, Ilia Gimaev, Michaela Fehringer, Márcia S. Pereira, Stefanie Robeck, Juheon Chu, Melanie Hagen, Andreas Wirsching, Tobias Rothe, Melanie Prinzensteiner, Aline Bozec, Fabian Müller, Andreas Mackensen, Thomas Vogl, Georg Schett, Ilana L. Brito, Martin A. Kriegel, Mario M. Zaiss

## Abstract

CD19-targeted CAR T-cell therapy induces profound B-cell depletion and clinical responses in autoimmune disease; however, whether it reshapes host–microbiota interactions remains unknown. We longitudinally profiled gut microbiome composition and microbiota-specific humoral immunity in 20 patients with systemic lupus erythematosus, systemic sclerosis, or idiopathic inflammatory myopathy undergoing CAR T-cell therapy, alongside 23 healthy controls, using shotgun metagenomics, IgA/IgG-bound microbiota profiling, and PhIP-Seq. CAR T-cell therapy reshaped microbiota-directed humoral immunity, with reduced fecal IgA and serum anti-commensal IgA and IgG accompanied by selective remodeling of microbiota-reactive antibody repertoires. This finding was paralleled by selective changes in gut microbial composition, including depletion of *Bifidobacterium adolescentis,* without uniform depletion of disease-associated pathobionts. Integrated host–microbiome analyses identified altered predicted protein-protein interactions involving the immunoproteasome and B-cell/plasma-cell pathways, with interactions before CAR-T cell intervention being significantly associated with subsequent relapse. Consistent with these findings, taxon-specific IgA targeting, but not global diversity of IgA/IgG-coated microbiota, differed according to subsequent relapse. Lower baseline fecal IgA was also associated with delayed B-cell reconstitution, linking mucosal humoral immunity to immune recovery after CAR T-cell therapy.

## Main Text

B cells are central drivers of autoimmune diseases (AIDs) through autoantibody production, antigen presentation, and cytokine production. Although targeted therapies have improved disease control in AIDs, these treatments suppress rather than reset autoimmunity and therefore require long-term treatment. Autologous CD19-directed chimeric antigen receptor (CAR) T-cell therapy provides a unique opportunity to study immune resetting by inducing profound B cell depletion followed by immune reconstitution. Sustained drug-free remission after a single CAR T-cell infusion has been reported in patients with severe AIDs, including systemic lupus erythematosus (SLE), systemic sclerosis (SSc), and idiopathic inflammatory myopathies (IIMs) ^1–3^. However, the biological processes accompanying immune reconstitution and durable remission after CAR T-cell-mediated immune reset remain poorly understood.

The intestinal microbiota is increasingly recognized as an important regulator of systemic immunity. Microbial antigens, metabolites, and host–microbial interactions can shape innate and adaptive immune responses, while microbiota-derived short-chain fatty acids (SCFAs) ^4–6^ and indoles ^5^ or inosine ^7^, as well as the host-derived ketone body β-hydroxybutyrate, can influence CAR T-cell fitness and persistence through metabolic and epigenetic pathways ^8^. Conversely, broad-spectrum antibiotic exposure around CAR T-cell infusion has been associated with inferior outcomes in hematological malignancies, and specific microbial communities and metabolites have been linked to responses to immune checkpoint blockade and CD19 CAR T-cell therapy in oncology ^9–11^. Whether these host–microbiota relationships are remodeled during CAR T-cell therapy in patients with AIDs and whether microbiome-associated changes are associated with the outcome of AIDs is currently unknown.

We longitudinally examined the gut microbiome and microbiota-directed humoral immunity before and after CAR T-cell therapy in 20 patients with refractory AID treated with CD19-targeting CAR T-cells within the CASTLE phase 1/2 basket study (NCT06347718), including patients with SLE (n = 9), SSc (n = 8), and IIM (n = 3) ^3^. We collected stool samples at baseline, the day before a single infusion of zorpocabtagene autoleucel (consisting of 1×10^6^ autologous CD19-CAR T-cells/kg body weight), and at 7 days, 2 months, and 6 months after the CAR T-cell infusion, along with samples from 23 age- and sex-matched healthy controls (Fig. 1a and Extended Data Table 1). In addition, blood samples were obtained from patients with AID (Fig. 1a). As previously reported, CD19-CAR T-cell therapy induces profound B-cell depletion, followed by B-cell reconstitution and sustained clinical remission ^12^. With extended follow-up of the present cohort, disease relapses occurred in 25% of the patients. Consistent with deep B-cell depletion, total serum IgA and IgG levels were reduced after CAR T-cell treatment (Fig. 1b,c). Despite standardized input, PhIP-Seq profiling of the antibody repertoire revealed that antigen recognition patterns post–CAR T-cell therapy tend to diverge from baseline, indicating a long-lasting shift in repertoire composition (Fig. 1d,e).

**Fig. 1.**
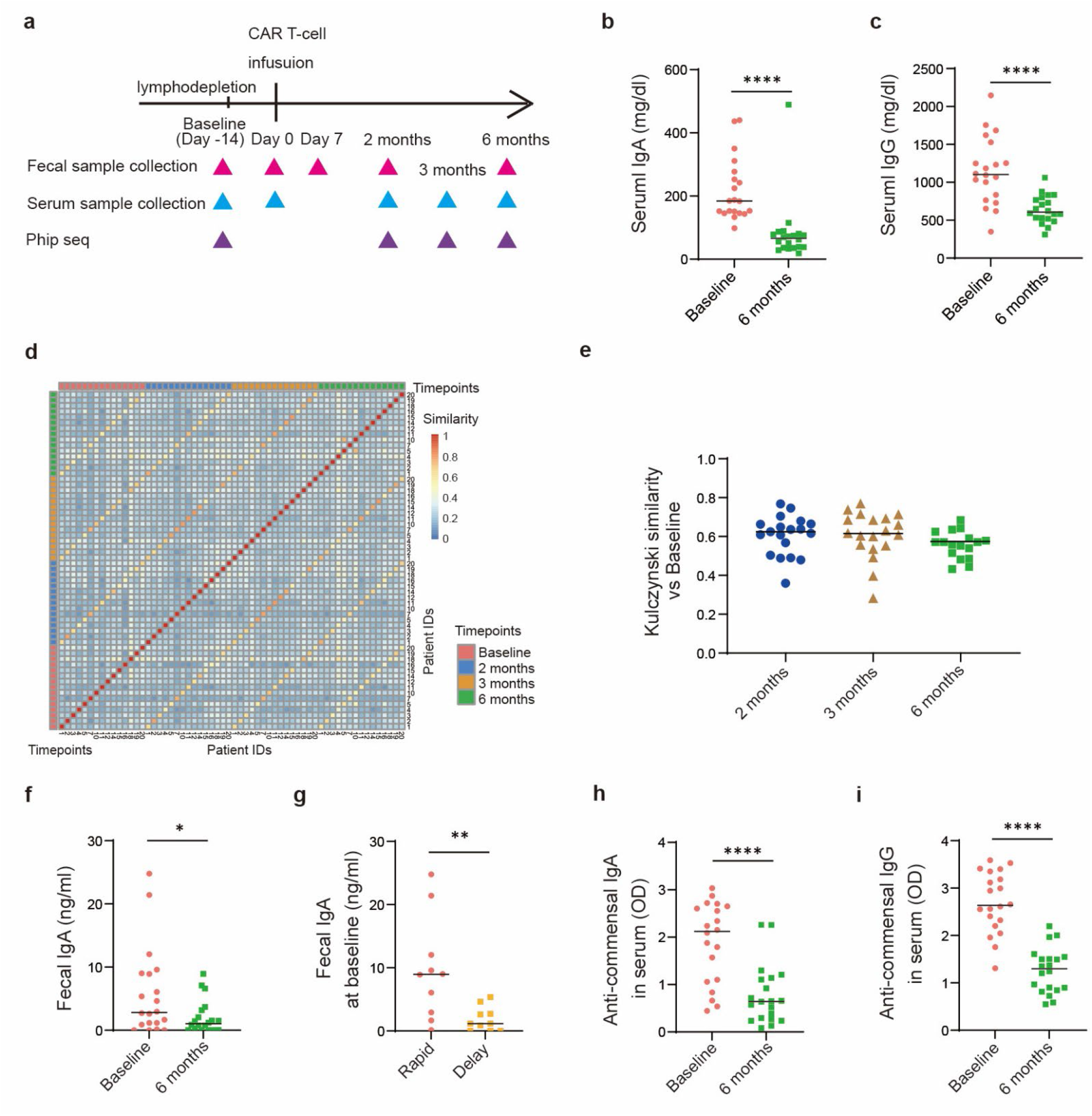
CAR-T cell therapy changes microbiota-directed humoral immunity. **a**, Study design and longitudinal sampling. **b**, Serum IgA and **c**, IgG concentrations at baseline and 6 months after the CAR-T cell therapy. **d, e**, PhIP-Seq results showing that the overall similarity of the antibody repertoire (quantified via Kulczynski similarity index) decreases compared to baseline. *P* values are shown as \**p* < 0.05, \*\**p* < 0.01, or \*\*\*\**p* < 0.0001. **f**, Fecal IgA concentration (baseline vs. 6 months). **g**, Fecal IgA concentration at baseline in patients with rapid B-cell recovery vs delayed B-cell recovery). **h, i**, Serum anti-commensal IgA and IgG levels (baseline vs 6 months).

We then assessed whether mucosal humoral immunity was similarly altered. Fecal IgA levels were reduced 6 months after CAR T-cell therapy compared to baseline (Fig. 1f). Given that peripheral B-cell recovery was delayed in a subset of patients (Extended Data Fig. 1a), we examined its association with baseline fecal IgA levels. Lower baseline fecal IgA levels were associated with delayed B-cell recovery (Fig. 1g and Extended Data Table 2). In contrast, serum immunoglobulin levels, microbiome diversity, and prior immunosuppressive treatment were not associated with delayed B-cell recovery (Extended Data Fig. 1b-e). Given the role of intestinal IgA in restricting systemic exposure to commensal microbes ^13,14^, we assessed the serum antibodies against commensal bacteria. Anti-commensal IgA and IgG levels were reduced after CAR T-cell therapy (Fig. 1h,i), while other antibody responses were largely preserved. Together, these findings suggest that CAR T-cell therapy depletes B cells and resets the host bacteria-specific humoral immune landscape.

Metagenomic profiling revealed a distinct gut microbial signature in patients with AID. Alpha diversity was reduced at baseline and further decreased after CAR T-cell therapy compared to healthy controls (Fig. 2a). Beta diversity based on Aitchison distance also differed according to disease status and CAR T-cell therapy, which together explained 7.8% of the variance in community composition (Fig. 2b). These global diversity measures did not differ between patients with and without relapse at either time point (Extended Data Fig. 2a,b). Patients with AID displayed a distinct gut microbial signature compared with healthy controls, characterized by enrichment of oral-associated taxa, including *Streptococcus* and *Actinomyces* species (Fig. 2c). Longitudinal analysis identified selective changes in microbial taxa following CAR T-cell therapy, with *Bifidobacterium adolescentis* showing the most pronounced depletion among abundant species (Fig. 2d), particularly in patients with SLE (Fig. 2e). Species-specific qPCR and anaerobic culture confirmed the reduction in detectable *B. adolescentis* following CAR T-cell therapy, consistent with loss of viable bacteria rather than changes in microbial DNA alone (Extended Data Fig. 3a,b). PhIP-Seq showed a trend towards reduced antibody reactivity to certain microbiota, including *B. adolescentis*, after therapy (Extended Data Fig. 3c). Previously described “AID-associated” pathobionts were not consistently depleted following CAR T-cell therapy. *Ruminococcus gnavus*, prevalent in patients with SLE at baseline, increased longitudinally after therapy, while *Enterococcus gallinarum* ^15^, detected only in a subset of patients, showed a similar increase, although a genomic resolution would be needed to determine if pathogenic strains were altered ^16^ (Extended Data Fig. 4a,b). *Lactobacillus reuteri*, which has interferonogenic potential ^17^, was not broadly prevalent in this cohort but showed a trend towards reduction in selected patients (Extended Data Fig. 4c). We also assessed circulating SCFA levels and fecal calprotectin levels but found no substantial changes following CAR T-cell therapy (Extended Data Fig. 5a,b). Thus, CAR T-cell therapy was associated with selective remodeling of the gut microbiome rather than consistent depletion of “AID-associated” pathobionts.

**Fig. 2.**
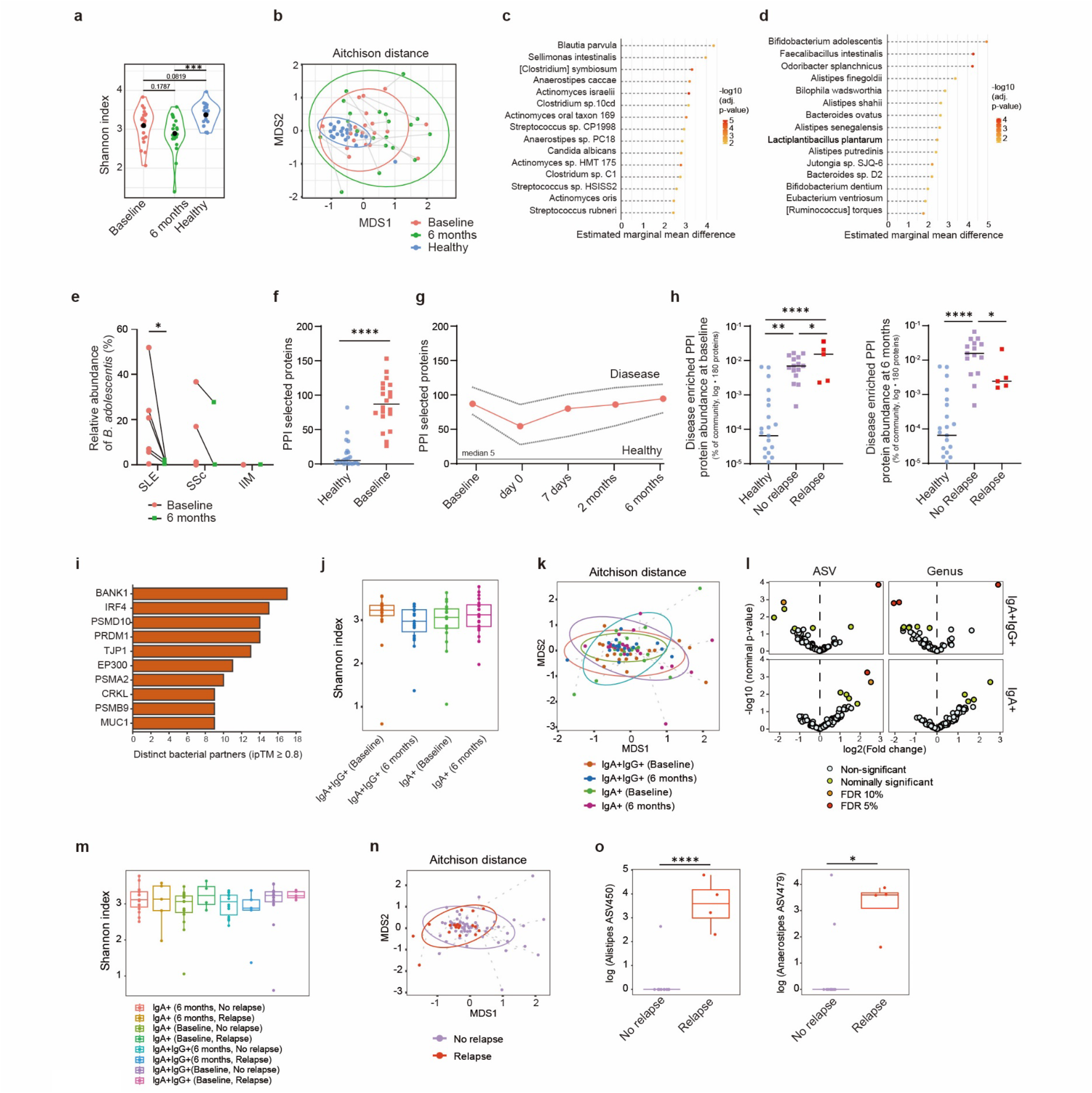
CAR-T cell therapy remodels the gut microbial landscape and host–microbial interactions. **a**, Metagenomic α-diversity by Shannon index (baseline vs. 6 months vs. healthy). **b**, Metagenomic β-diversity by Aitchison distance (baseline vs. 6 months vs. healthy). **c**, Top 15 taxa enriched before CAR T-cell therapy compared to healthy controls. **d**, Top 15 taxa reduced at 6 months after CAR T-cell therapy compared with baseline. **e**, Relative abundance of *Bifidobacterium adolescentis* in SLE, SSc and IIM (baseline vs 6 months). **f**, Protein–protein interaction (PPI) analysis of disease-enriched interactions at baseline (87 in patients vs 5 of 180 in healthy; Mann–Whitney U test, MWU, p=2e-07). **g**, PPI analysis of disease-enriched interactions across follow-up. **h**, PPI analysis at baseline and at 6 months (healthy vs. no-relapse vs. relapse). **i**, PPI analysis of the top 10 most targeted human proteins, ranked by the number of distinct bacterial interaction partners. **j**, IgG/IgA-seq α-diversity by Shannon index (baseline vs 6 months). **k**, IgG/IgA-seq β-diversity based on Aitchison distances at baseline and 6 months. **l**, Differential abundance analysis at the amplicon sequence variant (ASV) and genus levels was performed using linear mixed-effects models (IMER) and visualized as volcano plots. **m**, IgG/IgA-seq α-diversity by Shannon index (relapse vs no-relapse). **n**, IgG/IgA-seq β-diversity by Aitchison distance (relapse vs no-relapse). **o**, IgG/IgA-seq analysis showing significantly different ASV450 and ASV479 abundances (relapse vs no-relapse). *P* values are shown as \**p* < 0.05, \*\**p* < 0.01, \*\*\**p* < 0.001, or \*\*\*\**p* < 0.0001.

To examine host–microbiome interactions associated with microbial remodeling, we integrated metagenomic data with a host–microbiome protein–protein interaction (PPI) network analysis. Disease-enriched PPIs were more abundant in patients with AID than in healthy controls at baseline (Fig. 2f). They remained elevated throughout follow-up (Fig. 2g), despite lower microbial gene richness in patients with AID (median, ∼326,000 versus ∼377,000 UniRef90 gene families). Next, we examined whether host–microbiome PPI features were associated with subsequent relapse. At baseline, this measure was approximately 2.2-fold higher in patients who subsequently relapsed than in those who did not (MWU p = 0.35; relapse-label permutation p = 0.20, a patient-level test) (Fig. 2h and Extended Data Table 3). At 6 months after therapy, this abundance was approximately six-fold lower in patients who relapsed (0.16-fold; Mann–Whitney U test, p = 0.05; permutation test, p = 0.08) (Fig. 2h). Given the small relapse subgroup (n = 5), these longitudinal comparisons were considered to be exploratory.

The most pronounced PPI changes involved the immunoproteasome components PSMA2 (p = 4.70E-05) and PSMB9 (p = 2.48E-04) (Fig. 2i), which are associated with type I interferon (IFN-I)-driven inflammatory responses. As an IFN-I signature is a defining characteristic of AIDs^18^, these alterations may signify a universal trait. Several bacterial interactions with B cell-associated proteins were also among the most strongly altered PPIs, including BANK1 (p = 3.50E-11) and PRDM1 (also known as BLIMP1, p = 2.38E-08), which are key regulators of B cell signaling and plasma blast differentiation, respectively ^19^. Together, these findings link baseline host– microbiome PPI features to subsequent relapse and highlight altered interactions involving IFN-I-associated and B-cell/plasma-cell pathways following CAR T-cell therapy.

We next asked whether the reduced microbiota-specific serum IgA and IgG responses observed after CD19 CAR T-cell therapy (Fig. 1f, g) reflected a global loss of antibody reactivity or selective changes in microbial targeting. IgA- and IgG-Seq showed no significant changes in the α- or β-diversity of antibody-coated microbiota following CAR T-cell therapy (Fig. 2j,k). However, differential abundance analysis identified significant changes in IgA- and IgG-coated taxa at the genus and ASV level (Fig. 2l), indicating selective remodeling of antibody binding despite preserved overall diversity. ANCOM-BC analysis revealed a significant post-CAR-T decrease in *Prevotella* and the *Ruminococcus gnavus* group in the IgA⁺IgG⁻ subgroup (FDR-adjusted p < 0.05). In addition, decreases for *Bifidobacterium* and *Faecalibacterium*, albeit not significant, were recorded. We then asked whether these microbiota-directed antibody responses differed between patients who subsequently relapsed and those who remained in remission. Although alpha and beta diversity showed trends towards separation between groups, neither reached statistical significance (Fig. 2m,n). In contrast, differential abundance analysis identified distinct antibody-coated taxa associated with subsequent relapse. IgA-Seq revealed enrichment of specific IgA-bound microbial features, including *Alistipes* ASV450 and *Anaerostipes* ASV479, in patients who subsequently relapsed (Fig. 2o). *Prevotellaceae* UCG001 ASV145 also tended to increase in patients who subsequently relapsed (Extended Data Fig. 6). Thus, taxon-specific IgA binding, rather than global diversity of antibody-coated microbiota, distinguished patients according to subsequent disease relapse.

This study defines the longitudinal changes in the gut microbiome and mucosal humoral immunity during CD19-CAR T-cell therapy in AID. Patients with AID showed reduced microbial α-diversity and enrichment of oral-associated *Streptococcus* species^20^ at baseline, which are predicted to relocate from the oral cavity to the lower gastrointestinal tract. Following therapy, *Bifidobacterium adolescentis* was selectively depleted, particularly in patients with SLE. Given its reported capacity to induce Th17 -mediated inflammatory responses^21^, changes in this species may be relevant to immune remodeling. Notably, CD19-CAR T-cell therapy was accompanied by a remodeling of the host–microbiome immune interface, with reduced fecal IgA and microbiota-specific antibody responses, whereas pathogen-directed serological memory was relatively preserved. These findings are consistent with the differential depletion of CD19-expressing B cells, but not long-lived plasma cells, and suggest selective resetting of microbiota-directed humoral immunity. Lower baseline fecal IgA levels were also associated with delayed B cell recovery, linking the mucosal humoral status to systemic immune reconstitution.

Extended follow-up identified disease relapse in 25% of patients and enabled exploratory analyses of the features associated with the subsequent recurrence of disease. Host–microbiome PPI features involving immunoproteasome components, including PSMA2 and PSMB9, differed between patients with and without relapse. The association of PSMB9 and PSMA2 with relapse suggests remodeling of proteasome-mediated antigen processing during immune reconstitution. PSMB9, an interferon-inducible immunoproteasome subunit, may reflect renewed inflammatory immune activity accompanying the re-emergence of the disease. Given the small number of relapse cases, these findings require validation in larger cohorts in the future.

This study is limited by its rather small sample size and the limited statistical power for multi-omics and disease-specific analyses. Nonetheless, this cohort is very well characterized, as it is prospectively recruited according to a pre-defined study protocol^3^. A further limitation is the fact that relapses are infrequent in AID patients receiving CAR T-cells. Hence, the number of relapses that could be analyzed herein is small, and the comparison between relapsing and remission patients with respect to microbial changes should be seen as exploratory in nature. Larger longitudinal cohorts will be required to validate these findings and determine whether host– microbiome interactions contribute to the durability of immune resetting after CAR T-cell therapy.

Together, these findings provide, to our knowledge, the first integrated analysis of microbiome and host–microbiome interactions following CAR T-cell therapy in patients with AID and identify specific taxa and pathways that warrant further mechanistic investigation.

## Data availability

Data requests will be reviewed by the corresponding authors at the Medical University of Vienna, Austria; Cornell University, Ithaca, New York, United States; and Universitätsklinikum Erlangen (UKER), Erlangen, Germany. Patient-related data not included in this study were generated as part of clinical trials and may be subject to patient confidentiality. We will review requests for the stool sequence data. Any data and materials that can be shared will be released through a material transfer agreement. Raw ASV counts and annotations, as well as shotgun pathway counts and accompanying clinical annotations, are provided in the Supplementary Data.

## Acknowledgements

We thank all study participants for making this work possible. We thank PD. Dr. Stefan Wirtz and the Core Unit Microbiome Analysis Center Erlangen (MACE) at UKER for sequencing. We thank Prof. Gregor Fuhrmann at FAU Erlangen for the use of their flow cytometer (DFG grant no. 511550821). We thank H. Danzer and U. Leonhardt for supporting the study. We are grateful to technical support from B. Schulte and E. Mickholz and PCR validation from K. Stuhlträger and C. Brune at the University of Münster. Friedrich Alexander Universität Erlangen-Nürnberg initiated and initially sponsored the study. Sponsorship was transferred to Miltenyi Biomedicine GmbH in November 2024.

## Author contributions

Y.M., G.S., M.A.K., and M.M.Z. contributed to study conception and study design. Y.M., S.R., I.G., T.R., and M.M.Z. contributed to laboratory work. Y.M., Y.Y.L., S.R., V.P., I.G., M.F., M.S.P., S.Ro., J.C., M.H., A.W., T.V., I.L.B., M.A.K., and M.M.Z. contributed to data analysis. Y.M., V.P., M.F., S.R., S.Ro., Y.Y.L., J.C., and M.M.Z. contributed to results visualization. Y.M., G.S., M.A.K., and M.M.Z. contributed to writing the original manuscript. A.M. produced the CD19-CAR T-cells. M.H., A.W., T.R., A.B., F.M., G.S., and M.M.Z. contributed to resources.

## Use of AI tools

During the preparation of this manuscript, the authors used ChatGPT (OpenAI) and DeepL for language editing and translation. After using these tools, the authors reviewed and edited the content as needed and took full responsibility for the content of the manuscript.

## Methods

### Study design and participants

Patients with severe SLE, SSC, or IIM received a single infusion of an autologous lentiviral-based second-generation CD19-CAR T-cell product (zorpocabtagene autoleucel; MB-CART19.1; Miltenyi Biomedicine, Bergisch-Gladbach, Germany) at a dose of 1×10⁶ cells/kg body weight. CAR-T cells were generated via lentiviral transduction using a second-generation construct encoding a CD19-specific binder and a 4-1BB co-stimulatory domain. All patients were treated in the phase I/II CASTLE study (NCT06347718)^3^ approved by the ethics committee of Friedrich-Alexander-University Erlangen (22-168-Az). Written informed consent for publication was obtained from all patients whose data are presented in this manuscript.

### Collection of fecal samples

Stool samples were collected before lymphodepletion, defined as baseline, and 6 months after CAR T-cell therapy. In some of the patients, additional stool samples were collected before CAR-T cell infusion (day 0), and at 7 and 2 months after the infusion for the longitudinal analysis. These samples were collected using Gut Alive kits, which preserve bacteria under anaerobic conditions. Stool samples were stored at -80 °C until used for the experiment.

### Laboratory measurements

Levels of serum IgA, IgG, and common autoantibodies associated with AID were measured in the routine laboratory. Delay of B cell recovery was defined as failure of CD19⁺ cell counts to exceed 50 within 6 months after treatment.

### Evaluation of fecal IgA

Fecal IgA concentration was measured by ELISA, as previously described ^22^. Briefly, fecal samples were weighed and diluted 10-fold with PBS (w/v). The mixture was homogenized and then subjected to stepwise centrifugation at increasing force to remove debris. The resulting supernatant was diluted 100-fold, and the diluted samples were analyzed for IgA using the Human IgA Uncoated ELISA Kit (Invitrogen, cat #88-50600-88) according to the manufacturer’s instructions.

### Detection of AAPA-IgG and AAPA-IgA in serum and fecal samples

High-binding 96-well enzyme-linked immunosorbent assay (ELISA) plates (Nunc MaxiSorp, 442404, Thermo Fisher Scientific) were coated overnight at 4°C with acetylated ovalbumin (Ace– OVA; Sebia GmbH) at 20 μg/mL in phosphate-buffered saline (PBS; 50 μL/well). Plates were washed five times with PBS containing 0.05% Tween-20 (PBS-T; STS0200-250ML, Sigma-Aldrich) and blocked with 3% bovine serum albumin (BSA; A7030-100G, Sigma-Aldrich) in PBS (200 μL/well) for 1.5 h at room temperature. After five washes with PBS-T, serum samples diluted 1:10 in 1.5% BSA, or fecal samples (prepared as a 1:10 dilution of the original material, corresponding to a final dilution of 1:100 in 1.5% BSA) were added (50 μL/well) and incubated overnight at 4°C. Plates were then washed five times with PBS-T and incubated for 1 h at room temperature with horseradish peroxidase (HRP)-conjugated anti-human IgG (2040-05, SouthernBiotech; 1:10,000 in 1.5% BSA) or HRP-conjugated anti-human IgA (2050-05, SouthernBiotech; 1:5,000 in 1.5% BSA) (50 μL/well). After seven washes with PBS-T, tetramethylbenzidine (TMB; 00-4201-56, Thermo Fisher Scientific) substrate (50 μL/well) was added for color development. The reaction was terminated with a stop solution (SS04, Thermo Fisher Scientific) (50 μL/well). Absorbance was measured at 450 nm with reference wavelength correction at 620 nm using a Sunrise microplate absorbance reader (Tecan, Männedorf, Switzerland). Data were acquired using Magellan version 7.5 software (Tecan) and further processed using Microsoft Excel 2019 (Microsoft Corporation, Redmond, WA, USA).

### Detection of serum antibodies binding autologous fecal antigens by ELISA

High-binding 96-well ELISA plates (Nunc MaxiSorp, 442404, Thermo Fisher Scientific) were coated overnight at 4°C with fecal samples diluted 1:10 from the original material (corresponding to a final dilution of 1:100 in PBS) (50 μL/well). Plates were washed five times with PBS-T and blocked with 3% BSA in PBS (200 μL/well) for 1.5 h at room temperature. After five washes with PBS-T, matched serum samples from the same individuals were added at a dilution of 1:10 in 1.5% BSA (50 μL/well) and incubated overnight at 4°C. Plates were then washed five times with PBS-T and incubated for 1 h at room temperature with HRP-conjugated anti-human IgG (2040-05, SouthernBiotech; 1:10,000 in 1.5% BSA) or HRP-conjugated anti-human IgA (2050-05, SouthernBiotech; 1:5,000 in 1.5% BSA) (50 μL/well). After seven washes with PBS-T, TMB substrate (00-4201-56, Thermo Fisher Scientific) (50 μL/well) was added for color development, and the reaction was stopped with stop solution (SS04, Thermo Fisher Scientific) (50 μL/well). Absorbance was measured at 450 nm with reference wavelength correction at 620 nm using a Sunrise microplate absorbance reader (Tecan, Männedorf, Switzerland). Data were acquired using Magellan version 7.5 software (Tecan) and further processed using Microsoft Excel 2019 (Microsoft Corporation, Redmond, WA, USA).

### PhIP-Seq assay

Serum samples collected at baseline, day 56, day 84, and day 180 (6 months) post-CAR T-cell therapy were profiled using phage-display immunoprecipitation sequencing (PhIP-Seq). Assays were conducted as previously established by Vogl et al. 2021^23^, with the variation of using 2 µg of antibodies at a 4,000-fold phage coverage per library variant. We employed a combined phage library comprising the Agilent ^23^, Twist ^24^, and Corona ^25^ libraries, totaling 357,000 peptides, including human microbiota, pathogens, and environmental/dietary antigens. Sequencing was carried out on an Illumina NextSeq machine.

### PhIP-Seq raw data normalization

Single-end Illumina sequencing was performed on the samples and the input library (as a reference). Raw reads covering the 75 bp peptide-encoding barcodes were aligned to the reference barcodes (assigned to expressed peptides) using bowtie2 (v2.5.5) ^26^, tolerating a maximum of two mismatches. Read counts per peptide clone were quantified using samtools (v1.23) and awk.

Significantly enriched clones were identified by fitting a three-parameter zero-inflated generalized Poisson (ZIGP) distribution model as described previously ^27^. The ZIGP distribution parameters were then regressed based on the input count, using the scam (v1.2-19) library to fit a strictly increasing (parameters lambda and theta) or strictly decreasing (parameter pi) cubic spline. Peptides were considered enriched if their observed count corresponded to a p-value of <= 0.05 after Holm correction for multiple testing.

### PhIP-Seq data analysis

Normalized PhIP-Seq data was processed in R (v4.5.1) using tidyverse (v2.0.0) and pheatmap (v1.0.13). We quantified antibody repertoire overlap using the Kulczynski similarity coefficient via the vegan package (v2.7-2). Similarity was calculated as *Kulczynski similarity coefficient* = 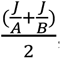, where J denotes the number of shared enriched epitopes, A and B represent the total number of bound epitopes identified in each sample.

### Flow cytometric detection of serum antibody binding to autologous fecal bacteria

An aliquot of 200 μL of the homogenized fecal suspension was centrifuged at 13000 rpm for 10 min. The resulting pellet was washed twice with 1 mL PBS, with centrifugation at 13000 rpm for 10 min after each wash. After removal of the supernatant, the pellet was resuspended in 100 μL of serum-PBS mixture (10 μL serum and 90 μL PBS) and incubated overnight at 4°C to allow serum antibodies to bind fecal bacteria. After incubation, 1 mL PBS was added, and the samples were centrifuged at 13,000 rpm for 10 min to remove unbound antibodies. The bacterial pellet was then incubated for 30 min at room temperature with Alexa Fluor 488-conjugated anti-human IgG antibody (M1310G05, BioLegend; 1:400 dilution in PBS) and Alexa Fluor 647-conjugated anti-human IgA antibody (2050-31, SouthernBiotech; 1:400 dilution in PBS) in a final staining volume of 100 μL. Samples were subsequently washed twice with 1 mL PBS, with centrifugation at 13000 rpm for 10 min after each wash, and finally resuspended in 200 μL PBS for flow cytometric analysis and cell sorting.

### Flow cytometric analysis and sorting of IgG- and IgA-coated bacteria

Flow cytometry analysis and fluorescence-activated cell sorting (FACS) were performed using a CytoFLEX SRT cell sorter (Beckman Coulter Life Sciences, Brea, CA, USA). Instrument acquisition, compensation, and sorting were controlled using CytExpert SRT software (Beckman Coulter Life Sciences; version 2.5). Flow cytometry data were analyzed using FlowJo software (version 10.10; BD Life Sciences, Ashland, OR, USA). IgG- and IgA-coated bacterial populations were identified based on the gating strategy and sorted for downstream analyses.

### DNA extraction from sorted bacterial populations

After sorting, the bacterial pellets were centrifuged at 13000 rpm for 10 min. After careful removal of the supernatant, the pellets were washed once with 1 mL PBS and centrifuged again under the same conditions. Genomic DNA was extracted using the QIAamp UCP DNA Micro Kit (Qiagen, Hilden, Germany) according to the manufacturer’s instructions. DNA concentration and purity were assessed using a NanoDrop 2000/2000c spectrophotometer (Thermo Fisher Scientific, Waltham, MA, USA) by measuring absorbance at 260 nm and evaluating the A260/A280 ratio. The extracted DNA was subsequently used for 16S rRNA gene sequencing.

### Extraction of bacterial DNA

Genomic DNA was extracted using the QIAamp PowerFecal Pro DNA Kit # 51804, according to the manufacturer’s instructions. These samples were analyzed for gut microbiota composition using metagenomic shotgun sequencing.

### Metagenomic shotgun sequencing and annotation

Fecal metagenomes were sequenced on the MiSeq platform (Illumina), including 20 patients from the CAR-T cell therapy for patients with autoimmune diseases longitudinally and 23 healthy controls. Shotgun sequencing raw reads were processed with fastp ^28,29^ (v1.0.1) and bowtie2 ^30^ (v2.5.4) against GRCh38 to remove low-quality reads and host contamination. Taxonomic annotation of the quality-filtered reads was performed using Kraken2 ^31^ and Bracken ^32^ at a 0.5 confidence threshold. Statistical analysis was performed in the R programming language v4.4.3 “Trophy Case” package. Alpha-diversity and beta-diversity were computed using the vegan R package (10.32614/CRAN.package.vegan). For alpha-diversity estimation, the Shannon index was used. For beta-diversity, the compositional-aware Aitchison metric was estimated; before the analysis, all singleton species were excluded, a pseudocount of one was added, and data were total- sum scaled. Projections of beta-diversity onto two-dimensional space were performed using the monoMDS function of the vegan R package. Alpha-diversity was modeled using mixed-effect linear regression implemented in the lme4 (10.32614/CRAN.package.lme4) and lmerTest R (10.32614/CRAN.package.lmerTest) packages. In equation (1), we included CAR-T status as the main independent variable, sex, age, DESeq2 ^33^ size factors, and scaled initial sequencing library size (*Seq_d*) as fixed covariates, and a patient-level varying intercept as a random effect.

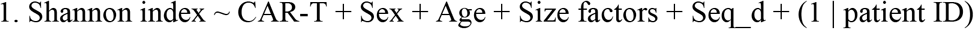

1. Shannon index ∼ CAR-T + Sex + Age + Size factors + Seq_d + (1 | patient ID)

Pairwise contrasts between groups were computed using the estimated marginal means approach implemented in the emmeans R package (10.32614/CRAN.package.emmeans) with Tukey’s correction for p-value adjustment. Beta-diversity was modelled using PERMANOVA for distance matrices with the adonis2 model of the vegan R package. The model (2) included a distance matrix (DM) as a dependent variable and the same set of covariates.

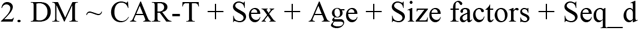

The model was run in marginal mode with 9,999 permutations. The homogeneity of within-group dispersions for CAR-T status was assessed using the betadisper function of the vegan R package with “type = median” and “bias.adjust = TRUE”. Differential abundance analysis was performed using mixed-effect linear regression implemented in the lme4 and lmerTest R packages. For the abundance of each species, we ran the regression equation (3) with a similar set of fixed and random covariates as for the alpha-diversity analysis. The significance of the three-group comparison was assessed using ANOVA type II, and estimated marginal means with FDR correction were used as a post-hoc test.

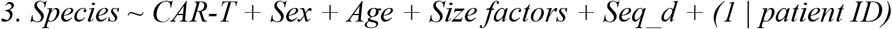

In addition to the mixed linear modelling, the ANCOM-BC2 package ^34^ was used for differential abundance analysis of 16S rRNA gene sequencing data. Before the analysis, read-based species abundance produced by Kraken2 was total-sum scaled and further transformed using a centered log-ratio transformation. Data were filtered to retain features present in at least 20% of samples in each group or in 50% of samples in any one group. In case of multiple comparisons, p-values were adjusted for multiple testing using the Benjamini and Hochberg procedure, and associations were considered significant at q-value < 0.05 after correction.

### Metagenomics profiling

Functional profiling was performed with HUMAnN3^35^ against UniRef90^36^ to obtain relative abundance of the UniRef90 gene-family table; we did taxonomic profiling using MetaPhlAn 3.1^35^ (database CHOCOPhlAn 201901).

### Selection of human target proteins

A panel of candidate human interactors was assembled based on their relevance to SLE (Extended Data Table 4). Candidates were collected from the following categories: (i) a manually curated list spanning interferon/inflammation signaling, TLR/NLR sensing, MHC class I and II, B- and plasma-cell identity, autoantibody target antigens, the core fibrosis program, intestinal barrier and mucin proteins, gut-homing receptors, and N-/O-glycosylation enzymes; (ii) the complete 26S proteasome, retrieved programmatically from the Gene Ontology^37,38^ through the QuickGO API^39^ as the union of the cellular-component terms GO:0005839 (proteasome core complex), GO:0008540 (proteasome regulatory particle, base subcomplex) and GO:0008541 (proteasome regulatory particle, lid subcomplex), including descendant terms over the ‘is_a’, ‘part_of’ and ‘occurs_in’ relations, restricted to Homo sapiens (NCBI taxon 9606); and (iii) the interferon-γ pathway gene set from the NCI-Nature Pathway Interaction Database^40^ as served by Harmonizome ^41^. Gene symbols were mapped to reviewed human UniProtKB accessions through the UniProt REST API^42^ (keyword: gene_exact + organism_id:9606 + reviewed:true). The panel comprised 265 gene symbols resolving to 264 unique accessions (P60896 has two different gene symbols), of which 262 human proteins entered the structural prediction analysis (MUC2 and MUC4 were excluded due to their large size).

### Selection of bacterial proteins

Candidate bacterial protein interactors were taken from a per-disease differential-abundance analysis of UniRef90 gene families, comparing each disease group at V1 against healthy controls (two-sided Mann-Whitney U test) on relative abundance with Benjamini-Hochberg correction. For each disease, we selected 250 significant bacterial proteins for the structural analysis.

### AlphaFold3 co-complex prediction

Each human protein was paired with each bacterial protein, and the complex was predicted with AlphaFold3 (AF3) v3.0.1 ^43^. Using AF3, we ran multiple sequence alignment (MSA) on Pittsburgh Supercomputing Center using Regular Memory compute nodes. Then we combine the paired and unpaired MSA from human and bacterial proteins, prepare the JSON file with five random model seeds. Then we run structural prediction on NCSA Delta using NVIDIA A100 GPUs. Of the 195,714 possible 262 × 747 combinations, 194,903 pairs were predicted. Confidence of co-complex prediction was scored by the interface-predicted template modeling score (ipTM) [doi:10.1101/2021.10.04.463034]. We filtered the predictions by ipTM of 0.8 or above, which are 542 pairs of putative protein-protein interactions, comprising 177 human and 212 bacterial proteins (where 180 of them are enriched in patients).

### Metagenomic readout of predicted interactions

For each sample, we combined the relative abundance of the bacterial gene families predicted to interact with the human proteins (ipTM of 0.8 or above) to demonstrate the potential combined effects targeting human proteins that play an important role in SLE. We restricted this to the 180 of 212 bacterial proteins enriched in patients at V1, and summarized each sample two ways: the number of those proteins detected, and summed abundance, their total relative abundance (% of community).

### 16S#rDNA-based bacterial species identification (single colonies)

Standard primers 16s_27f ^45^, 16s_1392r ^46^, 16s_1492r ^45^ were modified based on conserved regions in the 16S RNA gene: A database containing 120 representative 16S rDNA gene sequences was generated in accordance with ^47^ resulting in the modified primers 16s_27f_m (AGAGTTTGATCMTGGCTCAG), 16s_1392r_m (TGACGGGCGGTGTGTAC) and 16s_1492r_m (TACGTRTACCTTGTTACGAC).

Colony material of subcultures after separation was used directly for 16S rDNA PCR (16s_27f_m + 16s_1392r_m or 16s_27f_m + 16s_1492r_m). Resulting amplified DNA was purified (QIAquick PCR Purification Kit, Qiagen) and Sanger sequenced by an external provider (Eurofins). Sequences were checked with the BLASTn suite (BLAST, NCBI), restricted to rRNA databases and limited to type strain sequences.

### Quantitative PCR of bacterial taxa

DNA was extracted with the QIAamp PowerFecal Pro DNA Kit (Qiagen) following manufacturer’s instructions and quantified by UV-spectrometry (NanoDrop). Total bacterial 16S rRNA genes were quantified with universal primers (16s_uni_F_rt GATTAGATACCCTGGTAGTCC / 16s_uni_R_rt CGAGCTGACGACARCCATGC). Primers 16s_uni_F_rt and 16s_uni_R_rt were designed based on conserved regions in the 16S rDNA, amplifying a 268 bp core fragment of the gene. To identify conserved regions, a database containing 120 16S rDNA sequences was designed in accordance with^47^. *Bifidobacterium adolescentis* (BA) was quantified utilizing species-specific primers (BA-fwd GCTGATATCTGCGCTGTACC / BA-rev AAACCACCCAGTAGTCCTCC ^48^. Reactions (10 µL) contained 2 ng DNA (BA-specific vs. 16S rDNA), 0.25 µM of each primer, and 1 x SYBR Green master mix [LightCycler 480 Sybr Green I Master, Roche]. Samples were run in technical triplicates on a QuantStudio 5. 95 °C for 2 min, then 40 cycles of 95 °C for 15 s and 1 min at 56 °C (BA) or 58 °C (16S), followed by a melt curve (60–95 °C, 0.1 °C/s). Replicates were excluded if they showed no amplification, multiple melt peaks, or a Tm outside the assay window (BA 82.0– 83.8 °C, 16S 80.0–84.3 °C), or differed in their Ct by more than one cycle from the other replicates. Samples with values below the lowest standard were reported as below the limit of quantification. Samples without amplification in a valid reaction range were set to zero. Since the rDNA gene is present in multiple copies in bacteria, a correction factor was introduced. Calculated relative abundances of BA were multiplied by the factor (4.83). This factor was calculated based on species specific rDNA abundances per genome in representative microbiomes (mean ratio of rDNA per genome = 4.83; 0.275 standard deviation; n = 68). Final relative abundances are expressed as BA-specific gene copies per 16S gene copies (%). Correlation with metagenomics data was demonstrated by Spearman correlation.

### Fecal calprotectin by ELISA

Stool samples were collected with the GutAlive® Kit (Microviable Therapeutics, Spain) and aliquoted under anaerobic conditions. Fecal calprotectin was quantified from native stool aliquots using the ready-to-use ELISA kits from Immundiagnostik AG (Germany) “IDK® Calprotectin ELISA” and “Albumin ELISA” according to the manufacturer’s instructions.

