## Extended Figures 1-6 for "CD19-CAR T-cell therapy reshapes the host–microbiota immune axis in autoimmune diseases"

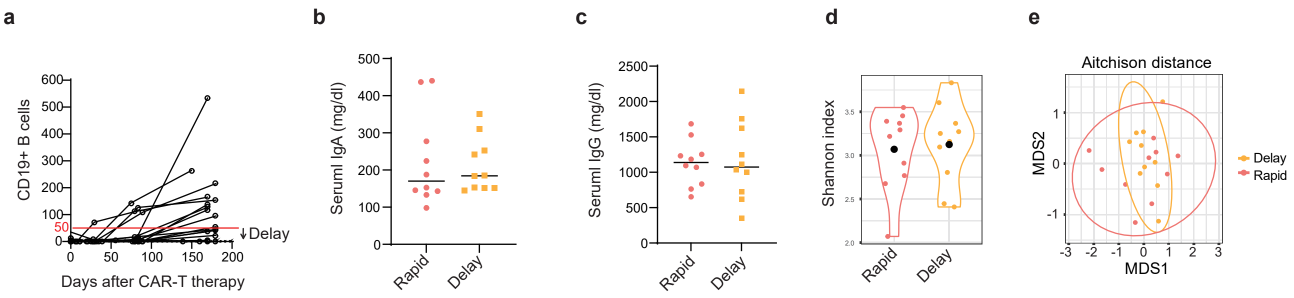


Extended Data Fig.1. Rapid or delayed B cell recovery after CAR-T cell therapy.

a, CD19+ B cell counts in the blood.

b, c, serum IgA (b) and IgG (c) concentrations at baseline (rapid vs. delayed B cell recovery).

d, e, Metagenomic α-diversity by Shannon index (d), β-diversity by Aitchison distance (e) at baseline (rapid vs. delayed B cell recovery).

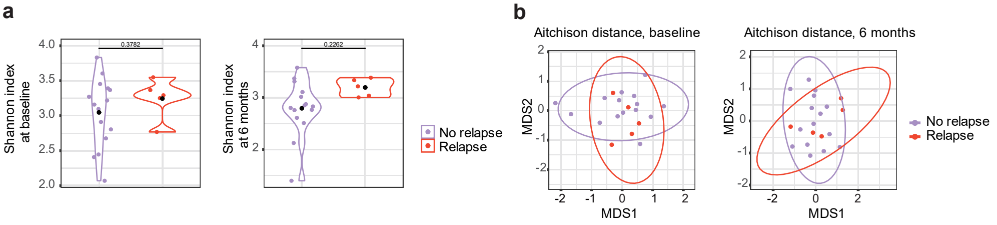


Extended Data Fig.2. α-and β-diversity of gut microbiota in patients with relapse or no relapse.

a, Metagenomic α-diversity by Shannon index at baseline and 6 months after CAR-T cell therapy (Relapse vs No relapse).

b, Metagenomic β-diversity by Aitchison distance at baseline and 6 months after CAR-T cell therapy (Relapse vs No relapse).

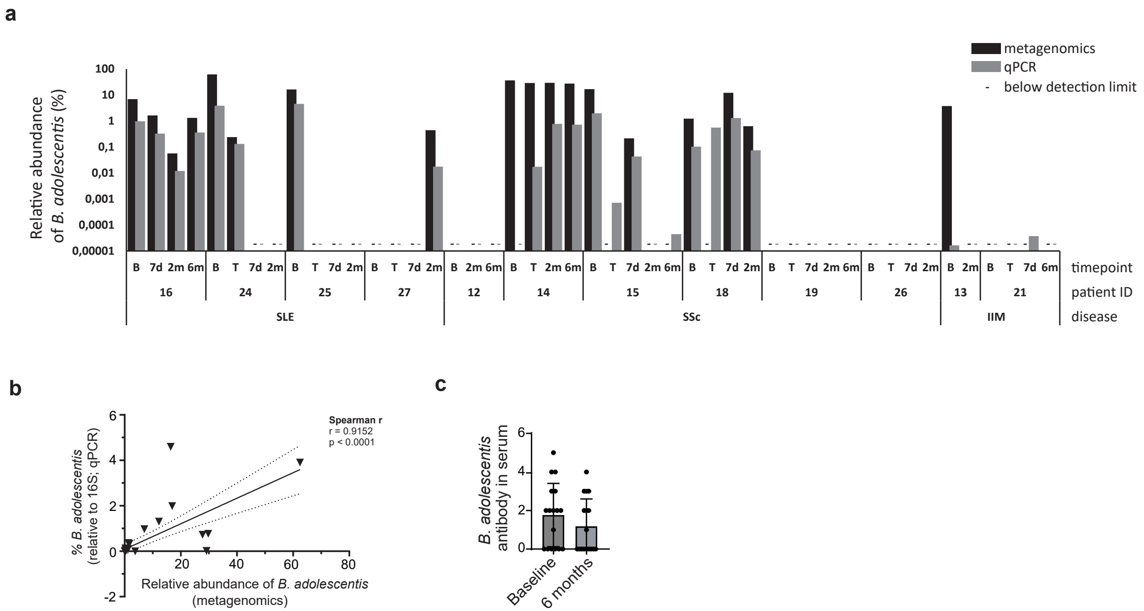


Extended Data Fig.3. Relative abundance of Bifidobacterium adolescentis.

a, Relative abundance of Bifidobacterium adolescentis measured by metagenomics and qPCR in patients with SLE, systemic sclerosis (SSc), inflammatory myositis (IIM). B (baseline), T (day 0), 7d (day 7), 2m (2 months), 6m (6 months).

b, Correlations between the B. adolescentis proportions measured by metagenomics and qPCR c, B. adolescentis antibodies in serum by PhIP-Seq (Baseline vs 6 months after CAR-T cell therapy).

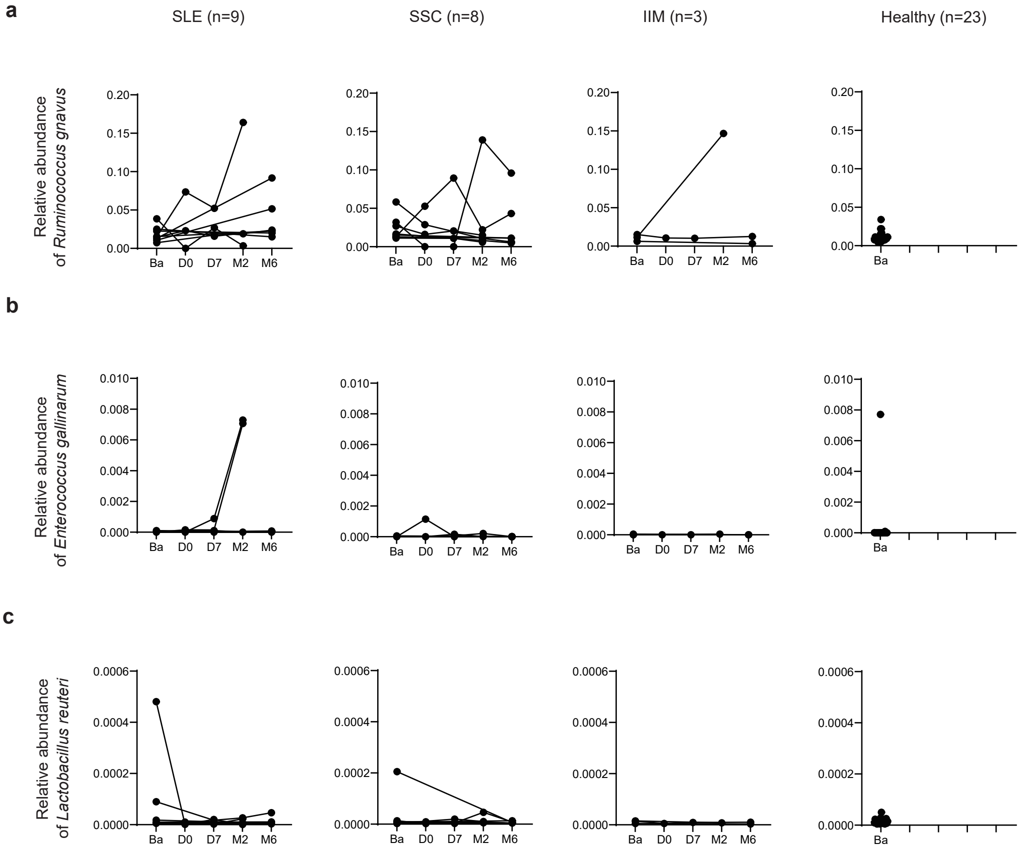


Extended Data Fig.4. Relative abundance of taxa previously associated with SLE.

a, Relative abundance of Ruminococcus gnavus.

b, Relative abundance of Enterococcus gallinarum.

c, relative abundance of Lactobacillus reuteri.

a

b

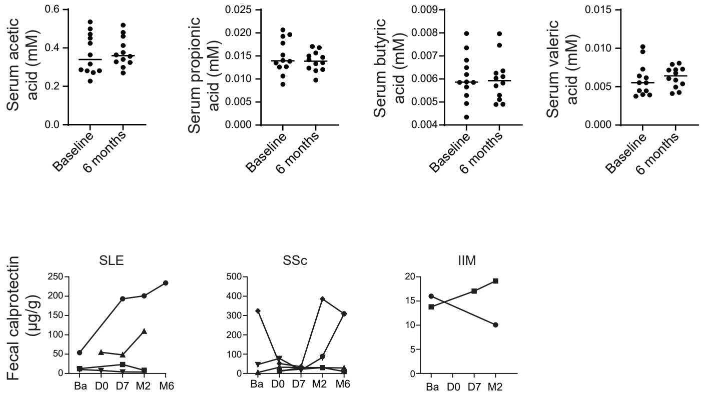


Extended Data Fig.5. Serum short-chain fatty acid (SCFA) and fecal calprotectin.

a, Serum short chain fatty acid levels (Baseline vs 6 months after CAR-T cell therapy). Serum acetic acid, propionic acid, butyric acid, and valeric acid levels are shown.

b, Fecal calprotectin levels in SLE, SSc, and IIM.

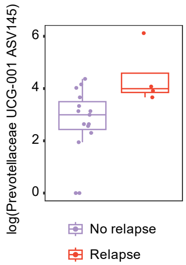


Extended Data Fig.6. Specific taxa identified by IgG/IgA-seq associated with relapse.

IgG/IgA-seq analysis showing differences in the abundance of Prevotellaceae UCG-001ASV145 (Relapse vs No-relapse).
