## Extended Data Tables 1-4 for "CD19-CAR T-cell therapy reshapes the host–microbiota immune axis in autoimmune diseases"

**Extended Data Table 1. Clinical characteristics of the patients who received CD19-CAR-T cell therapy, and healthy controls**

| Group | Patients (n=20) | | | Controls (n=23) |
|---|---|---|---|---|
| Disease | SLE (n=9) | SSc (n=8) | IIM (n=3) | |
| Age (years) | 33 ± 3.7 | 34.5 ± 5.3 | 56.0 ± 5.3 | 41.2 ± 12.9 |
| Sex, females (n) | 5 | 3 | 3 | 12 |
| Height (cm) | 170 ±3.1 | 176.5 ± 2.9 | 163 ± 2.1 | 172.5 ± 1.7 |
| Weight (kg) | 70 ± 4.2 | 71.5 ± 2.5 | 62.5 ± 1.1 | 78.2 ± 3.9 |
| Disease duration (years) | 5 ± 2.4 | 3.8 ± 0.4 | 1± 0.2 | N.T. |
| **Antibodies to (n)** | | | | |
| ANA | 9 | 8 | 2 | N.T. |
| Double stranded DNA | 9 | 0 | 0 | N.T. |
| Nucleosomes | 7 | 0 | 0 | N.T. |
| SSA/Ro60 | 5 | 0 | 0 | N.T. |
| Scl-70 | 0 | 7 | 0 | N.T. |
| RNA PolymeraseⅢ | 0 | 1 | 0 | N.T. |
| Ro-52 | 3 | 1 | 2 | N.T. |
| U1-RNP | 4 | 0 | 0 | N.T. |
| MDA5 | 0 | 0 | 1 | N.T. |
| Jo-1 | 0 | 0 | 1 | N.T. |
| **Organ involvement (n)** | | | | |
| Skin | 9 | 8 | 1 | 0 |
| Kidney | 7 | 0 | 0 | 0 |
| Gastrointestinal | 1 | 7 | 0 | 0 |
| Upper Gastrointestinal | 0 | 6 | 0 | 0 |
| Lower Gastrointestinal | 0 | 3 | 0 | 0 |
| Joints | 6 | 2 | 2 | 0 |
| Lung | 2 | 8 | 2 | 0 |
| Muscle | 0 | 4 | 3 | 0 |
| Haematological | 4 | 0 | 0 | 0 |

| | | | | |
|---|---|---|---|---|
| Heart | 3 | 4 | 0 | 0 |
| **Previous treatments** | | | | |
| Hydroxychloroquine | 8 | 0 | 0 | 0 |
| Azathioprine | 5 | 0 | 0 | 0 |
| Belimumab | 8 | 0 | 0 | 0 |
| Methotrexate | 6 | 6 | 0 | 0 |
| Cyclophosphamide | 2 | 1 | 0 | 0 |
| Mycophenolate mofetil | 6 | 7 | 1 | 0 |
| Prednisolone | 9 | 5 | 3 | 0 |
| Tocilizumab | 1 | 3 | 0 | 0 |
| Rituximab | 3 | 5 | 3 | 0 |
| Upadacitinib | 2 | 0 | 0 | 0 |
| Anifrolumab | 6 | 0 | 0 | 0 |
| Leflunomide | 2 | 0 | 0 | 0 |
| Obinutuzumab | 2 | 0 | 0 | 0 |
| Immunoglobulin | 0 | 1 | 2 | 0 |
| Previous antibiotics | | | | |
| Cotrimoxazole | 3 | 0 | 1 | 0 |
| Ciprofloxacin | 0 | 1 | 0 | 0 |
| Piperacillin/Tazobactam | 0 | 1 | 0 | 0 |
| Antibiotics after CAR-T | | | | |
| Cotrimoxazole | 8 | 8 | 3 | 0 |
| Ciprofloxacin | 6 | 2 | 1 | 0 |
| Piperacillin/Tazobactam | 5 | 4 | 1 | 0 |
| Meropenem | 3 | 2 | 0 | 0 |
| Vancomycin | 3 | 3 | 0 | 0 |

Unless otherwise noted, categorical variables are presented as numbers, and continuous variables as mean ± standard error (SE). ANA, antinuclear antibodies; anti-SSA/Ro60, anti–Sjogren's-syndrome-related antigen A/Ro60 autoantibodies; anti-SSA/Ro52, anti–Sjogren's-syndrome-related antigen A/Ro52 autoantibodies; Jo-1, anti–Jo-1 antibodies; MDA5, anti–melanoma differentiation-associated gene 5 antibody; RNA polymerase III, anti–RNA polymerase III antibody; Scl-70, anti–scleroderma 70 antibody; SLE, systemic lupus erythematosus; SSc, systemic sclerosis; IIM, idiopathic inflammatory myopathies; CAR, CD19 chimeric antigen receptor T cells; N.T., not tested.

**Extended Data Table 2. Baseline clinical characteristics of the patients with rapid versus delayed B-cell recovery.**

| **Group** | **Rapid recovery (n=10)** | **Delayed recovery (n=10)** | **P value** |
|---|---|---|---|
| Age (years) | 40.2±6.3 | 41.1±4.3 | 0.91 |
| Sex, females (n) | 6 | 5 | 1.0 |
| Height (cm) | 173.2±3.3 | 176±2.7 | 0.52 |
| Weight (kg) | 70.8±2.7 | 70.6±4.0 | 0.83 |
| Disease duration (years) | 2.5±0.6 | 7.3±2.4 | 0.08 |
| **Disease (n)** | | | - |
| SLE | 6 | 3 | 0.37 |
| SSc | 3 | 5 | 0.65 |
| IIM | 1 | 2 | 1 |
| **Antibodies to (n)** | | | |
| ANA | 9 | 10 | 1.0 |
| Double stranded DNA | 3 | 6 | 0.37 |
| Nucleosomes | 2 | 4 | 0.628 |
| SSA/Ro60 | 1 | 4 | 0.303 |
| Scl-70 | 4 | 3 | 1.0 |
| RNA PolymeraseⅢ | 1 | 0 | 1.0 |
| Ro-52 | 2 | 3 | 1.0 |
| U1-RNP | 2 | 2 | 1.0 |
| MDA5 | 0 | 1 | 1.0 |
| Jo-1 | 1 | 0 | 1.0 |
| **Previous treatments** | | | |
| Hydroxychloroquine | 5 | 5 | 0.65 |
| Azathioprine | 3 | 2 | 1 |
| Belimumab | 5 | 5 | 0.65 |
| Methotrexate | 6 | 6 | 1 |
| Cyclophosphamide | 2 | 1 | 1 |
| Mycophenolate mofetil | 8 | 6 | 0.628 |

| | | | |
|---|---|---|---|
| Prednisolone | 10 | 7 | 0.211 |
| Tocilizumab | 3 | 1 | 0.582 |
| Rituximab | 5 | 6 | 1 |
| Upadacitinib | 1 | 1 | 1 |
| Anifrolumab | 5 | 1 | 0.141 |
| Leflunomide | 1 | 1 | 1 |
| Obinutuzumab | 1 | 1 | 1 |
| Immunoglobulin | 3 | 0 | 0.211 |
| Previous antibiotics | | | |
| Cotrimoxazole | 3 | 1 | 0.582 |
| Ciprofloxacin | 0 | 1 | 1.0 |
| Piperacillin/Tazobactam | 1 | 0 | 1.0 |

Unless otherwise noted, categorical variables are presented as numbers, and continuous variables as mean ± standard error (SE). ANA, antinuclear antibodies; anti-SSA/Ro60, anti–Sjogren's-syndrome-related antigen A/Ro60 autoantibodies; anti-SSA/Ro52, anti–Sjogren's-syndrome-related antigen A/Ro52 autoantibodies; Jo-1, anti–Jo-1 antibodies; MDA5, anti–melanoma differentiation-associated gene 5 antibody; RNA polymerase III, anti–RNA polymerase III antibody; Scl-70, anti–scleroderma 70 antibody; SLE, systemic lupus erythematosus; SSc, systemic sclerosis; IIM, idiopathic inflammatory myopathies; CAR, CD19 chimeric antigen receptor T cells; N.T., not tested.

Continuous variables were compared between the two groups using Welch's t-test. Categorical variables, including the presence or absence of autoantibodies and medication use, were compared using Fisher's exact test.

**Extended Data Table 3. Baseline clinical characteristics of the patients with versus without relapse.**

| Group | Relapse (n=5) | No relapse (n=15) | P value |
|---|---|---|---|
| Age (years) | 45.0±7.0 | 39.1±4.4 | 0.46 |
| Sex, female (n) | 2 | 9 | 0.63 |
| Height (cm) | 179.0±3.3 | 173.1±2.5 | 0.19 |
| Weight (kg) | 77.0±2.5 | 68.4±2.7 | 0.06 |
| Disease duration (years) | 4.0±2.6 | 5.2±1.6 | 0.69 |
| **Disease (n)** | | | - |
| SLE | 1 | 8 | 0.319 |
| SSc | 3 | 5 | 0.347 |
| IIM | 1 | 2 | 1.0 |
| **Antibodies to (n)** | | | |
| ANA | 5 | 14 | 1.0 |
| Double stranded DNA | 1 | 8 | 0.319 |
| Nucleosomes | 0 | 7 | 0.114 |
| SSA/Ro60 | 1 | 4 | 1.0 |
| Scl-70 | 3 | 4 | 0.29 |
| RNA PolymeraseⅢ | 0 | 1 | 1.0 |
| Ro-52 | 3 | 3 | 0.131 |
| U1-RNP | 0 | 4 | 0.53 |
| MDA5 | 0 | 1 | 1.0 |
| Jo-1 | 1 | 0 | 0.25 |
| **Previous treatments** | | | |
| Hydroxychloroquine | 0 | 8 | 0.06 |
| Azathioprine | 1 | 4 | 1.0 |
| Belimumab | 1 | 7 | 0.603 |
| Methotrexate | 3 | 9 | 1.0 |
| Cyclophosphamide | 1 | 2 | 1.0 |
| Mycophenolate mofetil | 4 | 10 | 1.0 |

| | | | |
|---|---|---|---|
| Prednisolone | 4 | 13 | 1.0 |
| Tocilizumab | 1 | 3 | 1.0 |
| Rituximab | 2 | 9 | 0.617 |
| Upadacitinib | 0 | 2 | 1.0 |
| Anifrolumab | 1 | 5 | 1.0 |
| Leflunomide | 0 | 2 | 1.0 |
| Obinutuzumab | 0 | 2 | 1.0 |
| Immunoglobulin | 1 | 2 | 1.0 |
| Previous antibiotics | | | |
| Cotrimoxazole | 1 | 3 | 1.0 |
| Ciprofloxacin | 0 | 1 | 1.0 |
| Piperacillin/Tazobactam | 0 | 1 | 1.0 |

Unless otherwise noted, categorical variables are presented as numbers, and continuous variables as mean ± standard error (SE). ANA, antinuclear antibodies; anti-SSA/Ro60, anti–Sjogren's-syndrome-related antigen A/Ro60 autoantibodies; anti-SSA/Ro52, anti–Sjogren's-syndrome-related antigen A/Ro52 autoantibodies; Jo-1, anti–Jo-1 antibodies; MDA5, anti–melanoma differentiation-associated gene 5 antibody; RNA polymerase III, anti–RNA polymerase III antibody; Scl-70, anti–scleroderma 70 antibody; SLE, systemic lupus erythematosus; SSc, systemic sclerosis; IIM, idiopathic inflammatory myopathies; CAR, CD19 chimeric antigen receptor T cells; N.T., not tested.

Continuous variables were compared between the two groups using Welch's t-test. Categorical variables, including the presence or absence of autoantibodies and medication use, were compared using Fisher's exact test.

**Extended Data Table 4.** Selected panel of candidate human interactors was assembled based on their relevance to SLE .

| rank | uniprot_id | human_gene | n_distinct_bacterial_partners |
|---|---|---|---|
| 1 | Q8NDB2 | BANK1 | 17 |
| 2 | Q15306 | IRF4 | 15 |
| 3 | O75626 | PRDM1 | 14 |
| 4 | O75832 | PSMD10 | 14 |
| 5 | Q07157 | TJP1 | 13 |
| 6 | Q09472 | EP300 | 11 |
| 7 | P25787 | PSMA2 | 10 |
| 8 | P15941 | MUC1 | 9 |
| 9 | P28065 | PSMB9 | 9 |
| 10 | P46109 | CRKL | 9 |
| 11 | P05015 | IFNA16 | 8 |
| 12 | Q92793 | CREBBP | 8 |
| 13 | P25786 | PSMA1 | 7 |
| 14 | Q16584 | MAP3K11 | 7 |
| 15 | Q92985 | IRF7 | 7 |
| 16 | Q13761 | RUNX3 | 7 |
| 17 | P22681 | CBL | 7 |
| 18 | P12104 | FABP2 | 7 |
| 19 | P25789 | PSMA4 | 6 |
| 20 | O76094 | SRP72 | 6 |
| 21 | P01569 | IFNA5 | 6 |
| 22 | Q9H1M3 | DEFB129 | 6 |
| 23 | Q09327 | MGAT3 | 6 |
| 24 | P28072 | PSMB6 | 6 |
| 25 | P02461 | COL3A1 | 5 |
| 26 | P20618 | PSMB1 | 5 |
| 27 | P07199 | CENPB | 5 |
| 28 | Q30154 | HLA-DRB5 | 5 |
| 29 | Q9HBB8 | CDHR5 | 5 |
| 30 | Q13905 | RAPGEF1 | 4 |
| 31 | P28066 | PSMA5 | 4 |
| 32 | P49720 | PSMB3 | 4 |
| 33 | P28070 | PSMB4 | 4 |
| 34 | P10321 | HLA-C | 4 |
| 35 | P11473 | VDR | 4 |
| 36 | P17676 | CEBPB | 4 |
| 37 | P15391 | CD19 | 4 |
| 38 | P25788 | PSMA3 | 4 |
| 39 | P28062 | PSMB8 | 4 |
| 40 | O14818 | PSMA7 | 4 |
| 41 | A0A087WWS6 | IFNA13 | 4 |
| 42 | P01584 | IL1B | 4 |

| | | | |
|---|---|---|---|
| 43 | P01574 | IFNB1 | 4 |
| 44 | P02452 | COL1A1 | 4 |
| 45 | O15455 | TLR3 | 4 |
| 46 | O75925 | PIAS1 | 4 |
| 47 | Q14653 | IRF3 | 4 |
| 48 | Q13233 | MAP3K1 | 4 |
| 49 | Q00978 | IRF9 | 4 |
| 50 | Q8N2W9 | PIAS4 | 4 |
| 51 | Q9P0W0 | IFNK | 4 |
| 52 | Q86WN2 | IFNE | 4 |
| 53 | P05014 | IFNA4 | 3 |
| 54 | P01568 | IFNA21 | 3 |
| 55 | P05000 | IFNW1 | 3 |
| 56 | Q02556 | IRF8 | 3 |
| 57 | Q06141 | REG3A | 3 |
| 58 | Q13568 | IRF5 | 3 |
| 59 | Q9UMW8 | USP18 | 3 |
| 60 | Q53G44 | IFI44L | 3 |
| 61 | Q10981 | FUT2 | 3 |
| 62 | Q16186 | ADRM1 | 3 |
| 63 | Q14149 | MORC3 | 3 |
| 64 | Q14765 | STAT4 | 3 |
| 65 | Q16401 | PSMD5 | 3 |
| 66 | P32455 | GBP1 | 3 |
| 67 | P15291 | B4GALT1 | 3 |
| 68 | P62834 | RAP1A | 3 |
| 69 | P37173 | TGFBR2 | 3 |
| 70 | P17861 | XBP1 | 3 |
| 71 | P28074 | PSMB5 | 3 |
| 72 | P49721 | PSMB2 | 3 |
| 73 | P55036 | PSMD4 | 3 |
| 74 | P0DY56 | IFNA1 | 3 |
| 75 | O15263 | DEFB4A; DEFB4B | 2 |
| 76 | P01570 | IFNA14 | 2 |
| 77 | P01137 | TGFB1 | 2 |
| 78 | P09132 | SRP19 | 2 |
| 79 | P01889 | HLA-B | 2 |
| 80 | P08123 | COL1A2 | 2 |
| 81 | P01909 | HLA-DQA1 | 2 |
| 82 | Q9Y2R2 | PTPN22 | 2 |
| 83 | Q9UQM7 | CAMK2A | 2 |
| 84 | Q9UNM6 | PSMD13 | 2 |
| 85 | Q99436 | PSMB7 | 2 |
| 86 | Q8N687 | DEFB125 | 2 |
| 87 | Q8N104 | DEFB106A; DEFB106B | 2 |
| 88 | Q13557 | CAMK2D | 2 |
| 89 | Q13634 | CDH18 | 2 |

| | | | |
|---|---|---|---|
| 90 | Q12873 | CHD3 | 2 |
| 91 | Q02750 | MAP2K1 | 2 |
| 92 | P79483 | HLA-DRB3 | 2 |
| 93 | Q9H7Z7 | PTGES2 | 2 |
| 94 | Q9UPN9 | TRIM33 | 2 |
| 95 | Q6ZMB0 | B3GNT6 | 2 |
| 96 | P84022 | SMAD3 | 2 |
| 97 | Q13554 | CAMK2B | 2 |
| 98 | P61224 | RAP1B | 2 |
| 99 | P29466 | CASP1 | 2 |
| 100 | P31751 | AKT2 | 2 |
| 101 | P01571 | IFNA17 | 2 |
| 102 | P05013 | IFNA6 | 2 |
| 103 | O00231 | PSMD11 | 2 |
| 104 | O15105 | SMAD7 | 2 |
| 105 | O00487 | PSMD14 | 2 |
| 106 | P48556 | PSMD8 | 2 |
| 107 | P52630 | STAT2 | 2 |
| 108 | P23458 | JAK1 | 2 |
| 109 | P27986 | PIK3R1 | 2 |
| 110 | P11387 | TOP1 | 2 |
| 111 | P17181 | IFNAR1 | 2 |
| 112 | P15907 | ST6GAL1 | 2 |
| 113 | P10914 | IRF1 | 2 |
| 114 | P62191 | PSMC1 | 2 |
| 115 | P40306 | PSMB10 | 2 |
| 116 | P04439 | HLA-A | 1 |
| 117 | P08240 | SRPRA | 1 |
| 118 | P09914 | IFIT1 | 1 |
| 119 | P01911 | HLA-DRB1 | 1 |
| 120 | P01563 | IFNA2 | 1 |
| 121 | P01903 | HLA-DRA | 1 |
| 122 | P00973 | OAS1 | 1 |
| 123 | P01566 | IFNA10 | 1 |
| 124 | P01567 | IFNA7 | 1 |
| 125 | P01033 | TIMP1 | 1 |
| 126 | O14493 | CLDN4 | 1 |
| 127 | O15524 | SOCS1 | 1 |
| 128 | O15533 | TAPBP | 1 |
| 129 | O43242 | PSMD3 | 1 |
| 130 | O00206 | TLR4 | 1 |
| 131 | O14802 | POLR3A | 1 |
| 132 | O00233 | PSMD9 | 1 |
| 133 | P04083 | ANXA1 | 1 |
| 134 | P02778 | CXCL10 | 1 |
| 135 | P17706 | PTPN2 | 1 |
| 136 | P27361 | MAPK3 | 1 |

| | | | |
|---|---|---|---|
| 137 | P15260 | IFNGR1 | 1 |
| 138 | P26572 | MGAT1 | 1 |
| 139 | P12838 | DEFA4 | 1 |
| 140 | P13612 | ITGA4 | 1 |
| 141 | P19320 | VCAM1 | 1 |
| 142 | P18428 | LBP | 1 |
| 143 | P18084 | ITGB5 | 1 |
| 144 | P37108 | SRP14 | 1 |
| 145 | P40225 | THPO | 1 |
| 146 | P61626 | LYZ | 1 |
| 147 | P61011 | SRP54 | 1 |
| 148 | P59861 | DEFB131A | 1 |
| 149 | P60896 | SEM1 | 1 |
| 150 | P48551 | IFNAR2 | 1 |
| 151 | P42345 | MTOR | 1 |
| 152 | P42224 | STAT1 | 1 |
| 153 | P49450 | CENPA | 1 |
| 154 | P31749 | AKT1 | 1 |
| 155 | P29279 | CCN2 | 1 |
| 156 | P32456 | GBP2 | 1 |
| 157 | P32881 | IFNA8 | 1 |
| 158 | P35998 | PSMC2 | 1 |
| 159 | P35869 | AHR | 1 |
| 160 | Q09328 | MGAT5 | 1 |
| 161 | Q05655 | PRKCD | 1 |
| 162 | Q14116 | IL18 | 1 |
| 163 | Q13478 | IL18R1 | 1 |
| 164 | Q13555 | CAMK2G | 1 |
| 165 | Q07325 | CXCL9 | 1 |
| 166 | Q30KQ5 | DEFB115 | 1 |
| 167 | Q16625 | OCLN | 1 |
| 168 | Q8N688 | DEFB123 | 1 |
| 169 | Q7Z7B8 | DEFB128 | 1 |
| 170 | Q8NG35 | DEFB105A; DEFB105B | 1 |
| 171 | Q8TCB0 | IFI44 | 1 |
| 172 | Q9BYX4 | IFIH1 | 1 |
| 173 | Q96PH6 | DEFB118 | 1 |
| 174 | Q9BYC5 | FUT8 | 1 |
| 175 | Q9UHB9 | SRP68 | 1 |
| 176 | Q9Y275 | TNFSF13B | 1 |
| 177 | Q9Y5M8 | SRPRB | 1 |
